# Correlation of sympathetic and parasympathetic nervous system activity during rest and acute stress tasks

**DOI:** 10.1101/753657

**Authors:** David G. Weissman, Wendy Berry Mendes

**Author notes:** Corresponding Author: David G. Weissman, Harvard University, William James Hall, 33 Kirkland Street, Cambridge, MA, 02138.

## Abstract

Autonomic nervous system (ANS) activity is a core and central component of emotion. The myriad social and cognitive challenges faced by humans require flexible modulation of ANS activity for different contexts. In this study, simultaneous activity of the parasympathetic and sympathetic nervous system was measured using respiratory sinus arrhythmia (RSA) and pre-ejection period (PEP), respectively. Samples combined four previous studies (N=325) in which RSA and PEP were collected continuously during a resting baseline and an acute stressor, the Trier Social Stress Task. The concurrent relation between RSA and PEP responses was modeled in order to determine the extent to which SNS and PNS activity is correlated across the task within and between participants, and whether this correlation was moderated by age, race, sex, or baseline RSA and PEP. Overall, RSA and PEP were reciprocally coupled, perhaps reflecting shared regulatory mechanisms in the brain. However, recovery from a stressor was characterized by coactivation. Individuals also vary in the extent to which their SNS and PNS are reciprocally coupled; women, younger adults, and individuals with higher baseline RSA showed more reciprocal coupling than men, older adults, and those with lower baseline RSA, respectively, reflecting greater coordination of physiological responding in the former group.

## Introduction

Autonomic nervous system activity is a core and central component of the emotional response, although the centrality and specificity of autonomic responding as it relates to emotion and stress reactions are still topics of considerable debate and disagreement (Barrett, 2006; Friedman, 2010). Nonetheless, individual differences in autonomic responses to stress are valuable, unbiased indicators of vulnerability and resilience to mental and physical health problems (Alvares et al., 2016; Kemp & Quintana, 2013; Thayer et al., 2010). Autonomic activity can be measured by a variety of different measures, which, with varying degrees of mechanistic specificity, index the regulation of bodily systems in response to physical or mental tasks. Common notions of the function of the sympathetic nervous system (SNS) refer to it as the “fight or flight” system, suggesting that its activity contributes to high arousal, active states, whereas the parasympathetic nervous system (PNS) is referred to as the “rest and digest” system, suggesting an association between its activity and relaxation and restorative processes. However, the extent to which these two systems function on a single continuum or a more complex dynamic interplay is the subject of influential theorizing, and warrants systematic investigation.

### Autonomic Space

The SNS and PNS seemingly opposite roles may lead one to assume that their activity functions along a single-axis continuum. However, the autonomic space model (Berntson et al., 1991) suggests that the activity of the SNS and PNS are not universally reciprocal – when activity in one system increases activity in the other system always decreases. Instead, depending on the situation and individual differences, the two branches can shape peripheral responses, in particular heart rate (HR) through multiple forms of coordinated and uncoordinated activity, which can be represented as points in a two-dimensional state space (Figure 1). The possible activity patterns in this conceptualization include reciprocal patterns of activity, in which SNS and PNS activity are negatively correlated; SNS activation combined with PNS withdrawal leads to the reciprocal SNS state (top left) in which HR is elevated, while PNS activation combined with SNS withdrawal leads to the reciprocal PNS state (bottom right), in which HR is decreased. However, activity patterns can also be positively correlated, either through co-activation (top right) and co-inhibition (bottom left), leading to influences that counteract each other and therefore smaller changes in HR. Uncorrelated activity, through uncoupled increases or decreases in either system can also occur. The capacity to engage in different patterns activity across these two systems may allow for flexible modulation of somatic activity to meet diverse environmental demands. However, there are still gaps in our knowledge regarding how context and individual differences might influence these different relations between the two branches of ANS.

**Figure 1:**
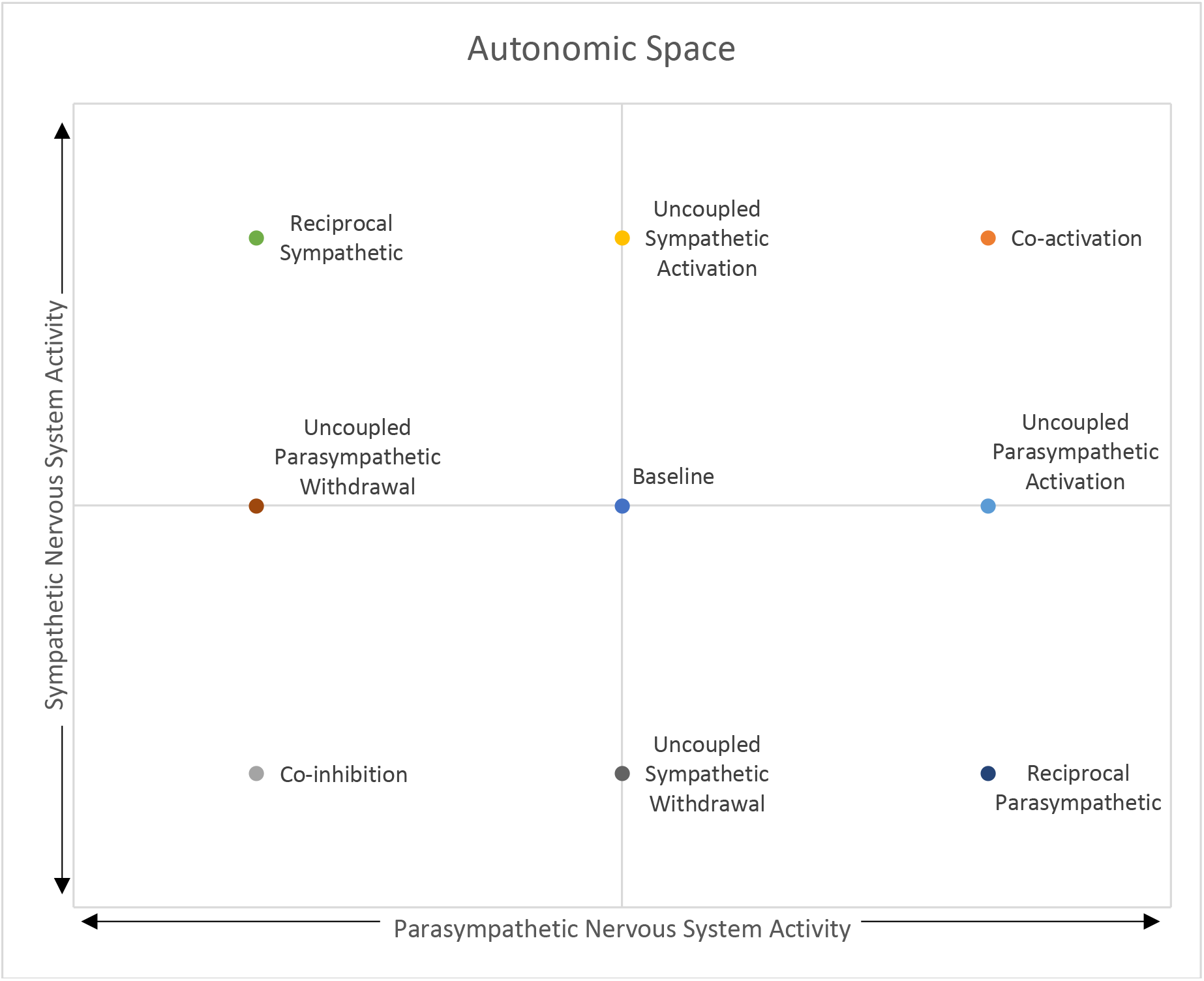
Theoretical representation of autonomic activity as a 2-dimensional state space. A representation of autonomic activity as a two-dimensional state space (Berntson et al., 1991)

The heart is under tonic inhibition by the vagus nerve. Thus, HR can decrease and increase by increases and decreases in PNS activity respectively. As would be expected by the law of initial values, higher baseline RSA is likely associated with greater capacity to increase HR through decreases in PNS activity (Berntson et al., 1991; Rigoni et al., 2017). Polyvagal theory argues that this higher baseline vagal activity impacts socioemotional functioning by allowing for more flexible emotional reactivity and regulation (Porges et al., 1994).

### Autonomic Responses to Stress

Autonomic responses are elicited when encountering and managing stressors (Allen & Crowell, 1989), an essential and unavoidable demand of daily life (Almeida, 2005) with consequences for emotion functioning and well-being (Serido et al., 2004). The Trier Social Stress Task (TSST) is a laboratory stressor designed to characterize an individual’s typical response to an acute social stressor in daily life (Kirschbaum et al., 2008). The TSST has robust, well documented effectiveness at inducing psychological distress and physiological stress responses (Allen et al., 2014), including a strong activation of the hypothalamic pituitary adrenal cortical axis (HPA) (Dickerson & Kemeny, 2004) Traditionally, the TSST involves the participant planning and delivering a short speech to stoic evaluators, followed by a mental arithmetic task, then, in some cases, a recovery period.

Measures of SNS and PNS influence on HR typically demonstrate consistent, reciprocal responses to social stress. Pre-ejection period (PEP) is a measure of the time between the electrical impulses that initiate ventricle contraction and the opening of the aortic valve, which is primarily influenced by the SNS. Decreases in PEP indicate an increase in SNS activity, at least in the absence of postural changes (Cacioppo et al., 1994a). Participants delivering evaluated speeches typically show and increase in heart rate and a decrease in PEP relative to baseline (Cacioppo et al., 1994a). Respiratory sinus arrhythmia (RSA) is high frequency heart rate variability in the respiratory frequency band that reliably estimates PNS control of the heart via the vagus nerve (Berntson et al., 1993). The effect of the TSST on PNS activity is less consistent across studies (Allen et al., 2014). However, a meta-analysis found a small but significant (*g*=−.27) effect size of decreases in RSA during the speech part of the TSST (Shahrestani et al., 2015), although it is possible that simply speaking itself in the absence of social stress or evaluation may alter RSA (Cacioppo et al., 1994b).

In one study, in which RSA and PEP were measured simultaneously during the TSST in a sample of adult women, the between-subjects correlations of RSA and PEP with HR were negative and positive, respectively, while the correlation of RSA with PEP was low to moderate and not significant in a relatively small sample size (Cacioppo et al., 1994b). The lack of a strong between-subjects relation between RSA and PEP reactivity during social stress suggests that the relative magnitude of SNS and PNS influence on the heart might be variable between people. Further, while evidence from pharmacological blockade of the SNS and PNS in the same sample suggests that the influence of the SNS and PNS on heart period responses to orthostatic stress (i.e. standing vs. sitting) are reciprocal, their influence on heart period responses to the TSST is uncorrelated (Cacioppo et al., 1994a, 1994b). Cross-situational, within person analyses of noninvasive measures of SNS and PNS functioning (PEP and RSA), such as through mixed effects modeling, can be revelatory in evaluating the nature of correlated activity between the two systems as individuals navigate different situations.

### Individual differences in autonomic responses to stress

Higher baseline PNS activity leads to a greater potential range of activation within which to modulate PNS responding to meet environmental demands. In response to acute stressors, individuals with higher baseline RSA suppress RSA to a greater extent. However, the magnitude of the recovery immediately following the stressor, as indicated by the difference between RSA during the stressor and RSA during a subsequent relaxation or recovery period, might not be associated with baseline levels of RSA (Rigoni et al., 2017). Individual differences along trait dimensions show that people who endorse more of a “belief in a just world” tend to show more SNS activation to a stressor (Tomaka & Blascovich, 1994), as do participants higher in self-esteem assessments (Seery et al., 2004). Older adults have been found to have larger decreases in both RSA (Uchino, Uno, Holt-Lunstad, & Flinders, 1999; Uchino, Holt-Lunstad, Bloor, & Campo, 2005) and PEP (Uchino et al., 1999) during laboratory stressors than younger adults, indicating a reciprocal autonomic response to stress with increasing magnitude of both SNS and PNS activity with increasing age. Females have been found to have higher resting heart rate variability (e.g. RSA) than males, despite having higher heart rates (Koenig & Thayer, 2016). Racial differences in PNS activity have also been observed, with black adult participants having higher baseline RSA than white participants (Dorr et al., 2007; Kemp et al., 2016). However, aggregate levels of activity may not capture variability in dynamic responses across multiple systems. To our knowledge, the nature of correlated responses in the PNS and SNS simultaneously has not been compared systematically as a function of participant age, sex, race, or baseline autonomic activity. By characterizing the nature of correlated PNS and SNS responses and how these differ based on sex, age, and race, we can better understand how stress responses are coordinated across the autonomic nervous system, paving the way for a better understanding of individual differences in emotion functioning and mental, and physical health.

### The Current Study

The current study combines data from four studies utilizing the TSST among adolescent and adult participants, diverse in age, race, and sex. In all studies, RSA and PEP were measured and data were binned in 1-minute increments during the five periods of the TSST 1) baseline, 2) speech prep, 3) speech, 4) mental math or question and answer (Q&A), and 5) recovery. A total of 25 minutes of physiological data was examined for each subject. The resulting large sample size allows us to use multilevel modelling approaches to examine both within and between-subject effects on correlated autonomic activity. This study aims to test the following hypotheses. 1) We expect that, overall, changes in RSA and PEP will be reciprocally coupled (i.e. positively correlated) during the TSST within individuals; 2) This pattern will vary based on the demands of the task, however. When the task is novel and activation is strongest – during the preparation and speech delivery time—reciprocal SNS activity is expected, whereas during the recovery period coactivation is expected, during what has been referred to as the “vagal rebound” (Mezzacappa et al., 2001; Page-Gould et al., 2010); 3) Given that higher baseline RSA is associated with a greater capacity to decrease RSA, we hypothesize that participants with higher baseline RSA will exhibit greater decreases RSA to accompany increases in PEP in response to stressors, which will be reflected in more reciprocal activity across the task. Whether individual differences in patterns of coupling are related to participant sex, race, and age, and whether those differences are independent of or mediated by baseline RSA was explored.

## Method

### Participants and Procedure

This study is a reanalysis of data from four previous studies where RSA and PEP were collected during the TSST. Exact procedures and details for each individual study are described elsewhere (Akinola & Mendes, 2008; Ayduk et al., 2012; Mendes & Koslov, 2013; Page-Gould et al., 2014). A total of 325 adult participants completed the studies. For all four studies, the TSST consisted of a 5-minute baseline, either a 3- or 5-minute speech preparation, followed by a 7- or 5-minute speech, and ending with a 5-minute recovery period. In all studies, participants remained seated for all task periods and were instructed to refrain from postural changes. Speech preparation was entirely mental. No writing was permitted. In two of the studies the speech was followed by mental arithmetic, and in the other two, it was followed by a question and answer task. In two of the studies, half of the participants received positive affective responses from the evaluators (they smiled and nodded during the TSST compared to the standard TSST evaluator feedback which includes the evaluators scowling and shaking their heads). These participants were included, but positive feedback was included as a covariate in all between-subject analyses. Sex was not recorded as a variable in one of the studies (N = 67). Age and race were only recorded as variables in one of the studies that focused on race differences. This study included 68 black and 73 white participants, 79 female and 62 male participants. Mean age was 29.0 (*SD* = 10.7, Range 15 to 55).

### Measures

#### Autonomic Nervous System

In all studies, electrocardiography (ECG100C) and impedance cardiography (HIC) were collected and integrated with an MP150 system (BIOPAC Systems Inc, Goleta, CA). Sensors to measure ECG were applied in a lead II configuration and impedance cardiography was obtained using four mylar bands that completely encircled the neck and chest area. A 1 mA AC current at 100 kHz was passed through the outer bands, and Z0 and dZ/dt were recorded from the inner bands. All signals were filtered on-line and sampled at 1000 Hz.

RSA was edited and analyzed using the HRV (2.5) module from Mindware Technologies (Gahanna, OH). Visual inspection of the waveforms focused on detecting ectopic beats and accurate detection of R spikes in the ECG. HRV was scored in 1-minute bins. The HRV module detrended the data using a first order polynomial to remove the mean and any linear trends, cosine tapered the data, submitted it to Fast Fourier Transformation, and took the natural log integral of the high frequency power (.15–.40 Hz) as an index of RSA.

PEP was edited and analyzed using the IMP (3.0) module from Mindware Technologies (Gahanna, OH) to edit impedance data. Visual inspection focused on accurate Q, R, S placement in the ECG trace and accurate detection of the B-, X-, and Z-points (aortic valve opening, aortic valve closing, and dz/dt max, respectively) on the dZ/dT waveform. PEP was calculated as the time interval in ms between the Q-point of the ECG and the B-point of the dZ/dt signal. PEP was averaged over 1-minute bins

### Analysis

Baseline RSA and PEP were be calculated by taking the mean of the 5 baseline bins. RSA reactivity (ΔRSA) and PEP reactivity (ΔPEP) within each of the task periods (including baseline) were calculated by subtracting baseline RSA and PEP from unstandardized RSA and PEP. Differences in the magnitude of ΔRSA and ΔPEP during each task period based on sex and race, and correlations with baseline RSA and PEP, age, and with the other indicator during the same task period were calculated.

Four two-level mixed effects structural equation model were tested in MPlus. This approach to identifying variability in the degree of temporal covariation between two physiological signals is best suited for data in which the number of physiological observations per subject (25) is less than the number of subjects (325) (Helm et al., 2018). In model 1, raw ΔRSA and ΔPEP were used in order to obtain meaningful and interpretable estimates of differences in RSA and PEP changes relative to baseline in each of the task periods. In models 2-4, ΔRSA and ΔPEP were standardized across the 25 1-minute epochs within subjects to normalize the variances of the two measures and obtain an unbiased measure of bidirectional synchrony.

Models 1 and 2 were tested across all 25 1-minute task epochs in all 325 participants. Model 1 tested the within-subjects relation between task period (prep, speech, math, Q&A, and recovery, relative to baseline) and ΔRSA, between task period and ΔPEP, and examine the residual covariance between ΔRSA and ΔPEP. Model 2 tested the within-subjects relation between ΔRSA and ΔPEP across all 25 1-minute epochs regardless of task period. At the between-subjects level, we tested whether the random slope of the within-subject association between ΔRSA and ΔPEP was moderated by baseline RSA, baseline PEP, the study that the data came from, or study conditions (i.e. whether the feedback subjects received from evaluators was positive or neutral).

Model 3 was estimated in the 258 participants whose sex was recorded. In addition to the parameters in Model 2, Model 3 tested whether sex moderated the within subject associations, and whether sex was associated with baseline RSA. The indirect effect of sex on the slope of the relation between ΔRSA and ΔPEP via baseline RSA were also estimated.

Model 4 was tested only among the 131 participants from the single study in which we had data on race and age. In addition to participant sex, baseline RSA, and baseline PEP, Model 4 also examine whether race and age moderated within subject associations between ΔRSA and ΔPEP. Paths were also included from sex, race, and age to baseline RSA and PEP. MPlus code for all analyses can be found at github.com/dgweissman/autonomic-balance.

## Results

### Mean autonomic activity

Mean baseline RSA was 6.74 (units = ln[ms^2^]; *SD* = 1.22). Mean baseline PEP was 118.4 (*SD* = 21.0). On average, participants’ RSA was significantly lower than baseline during prep, speech, math, and Q&A, and significantly higher than baseline during recovery. PEP was significantly shorter than baseline during prep, speech, math, Q&A, and recovery (Figure 2). Results are summarized within each of the 4 studies in Table 1. Figure 3 depicts these results plotted in a 2-dimensional state space.

**Figure 2:**
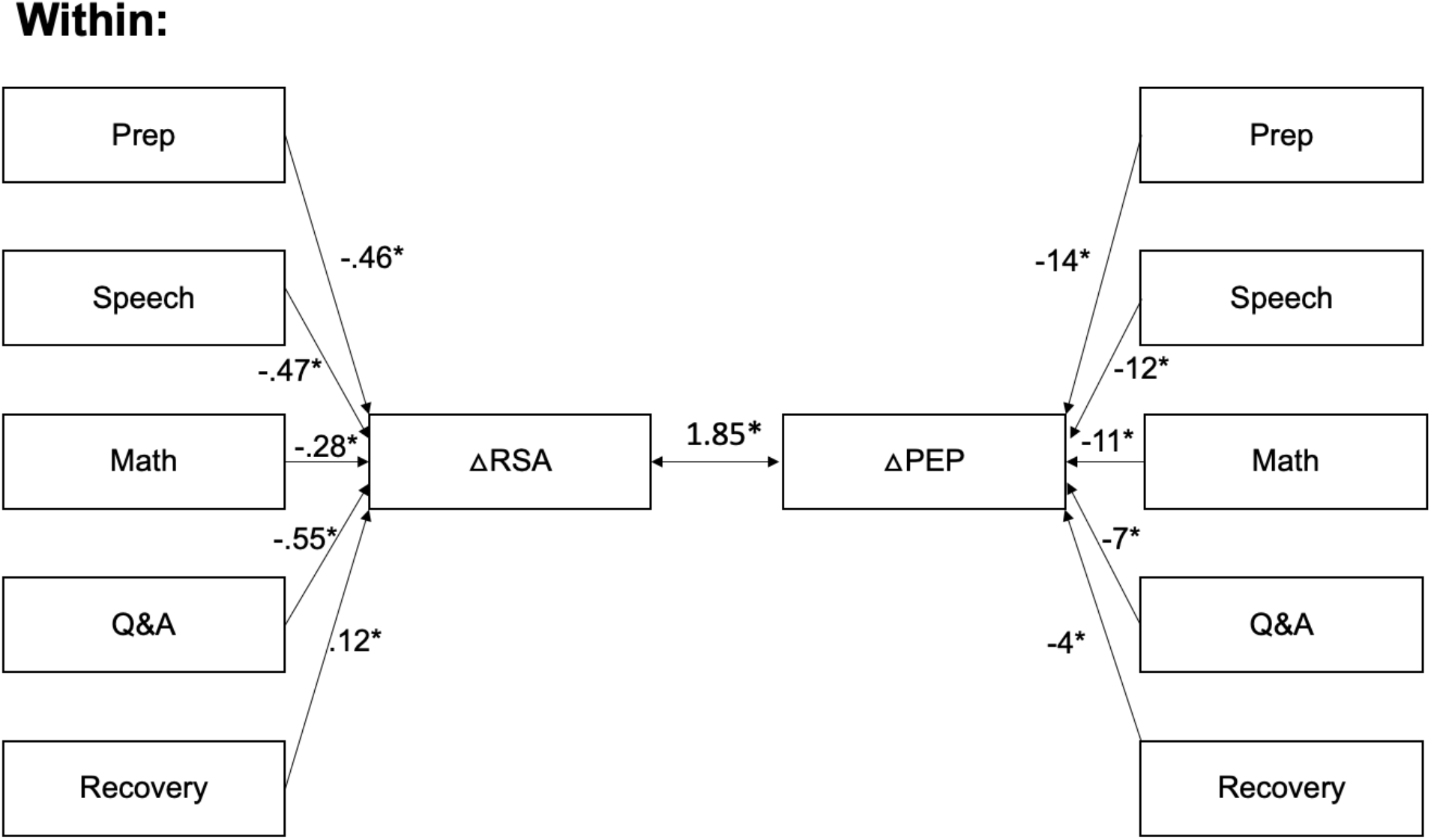
Within-person associations between task period and RSA and PEP and residual covariance. Results of multilevel structural equation modelling of factors contributing to within-person changes in RSA and PEP (N=325). The change in RSA (ΔRSA) and PEP (ΔPEP) relative to baseline was determined at each of the 25 1-minute epochs of the task. Coefficients represent the unstandardized change in RSA and PEP relative to baseline. The residual covariance is also unstandardized. RSA = respiratory sinus arrhythmia (ln[ms^2^]), PEP = pre-ejection period (milliseconds). * p < .05

**Figure 3:**
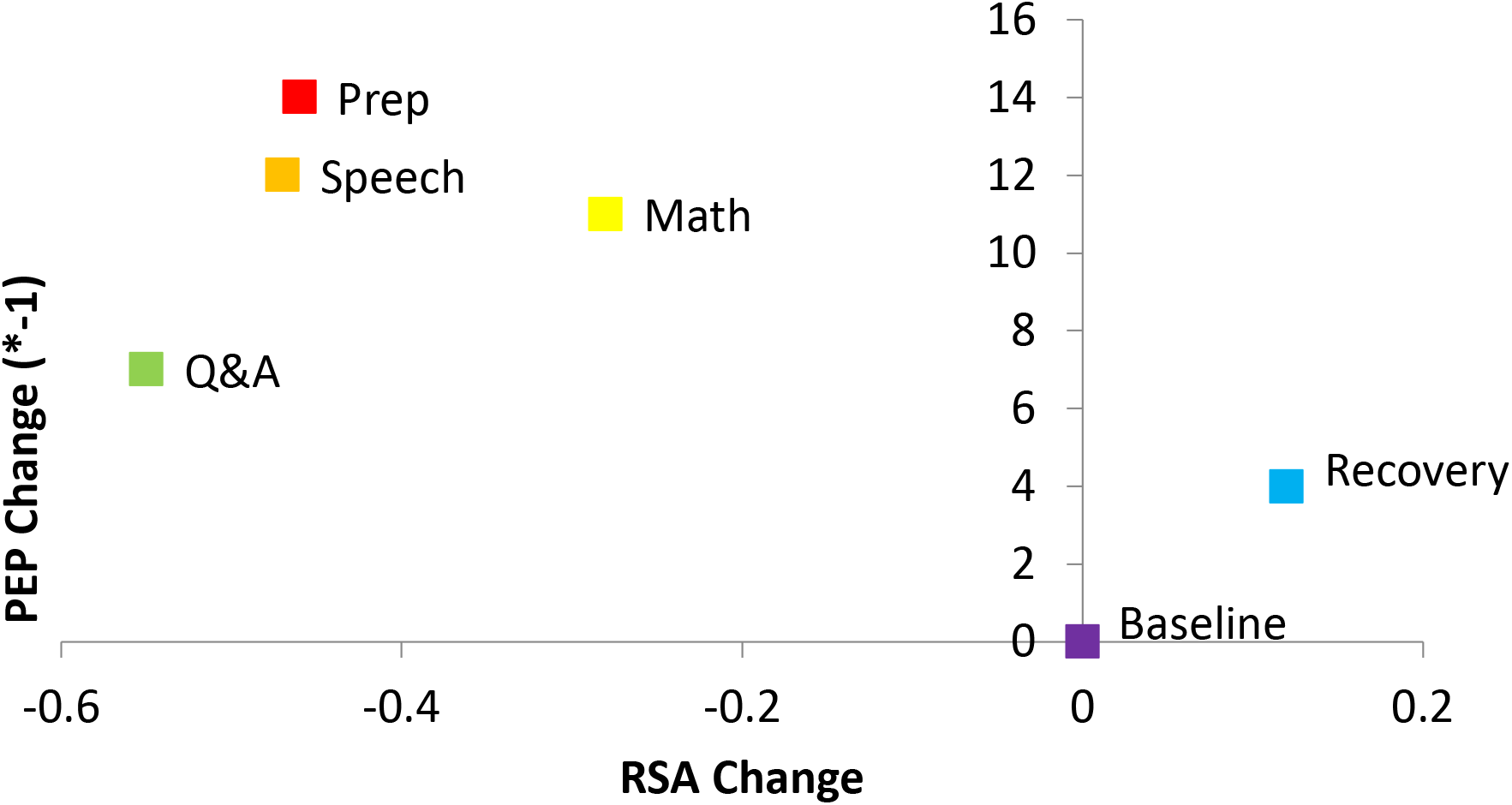
Change in autonomic activity during the Trier Social Stress Task. Mean change in autonomic activity during each part of the Trier Social Stress Task and recovery is plotted in a 2-dimensional state space.

**Table 1:**
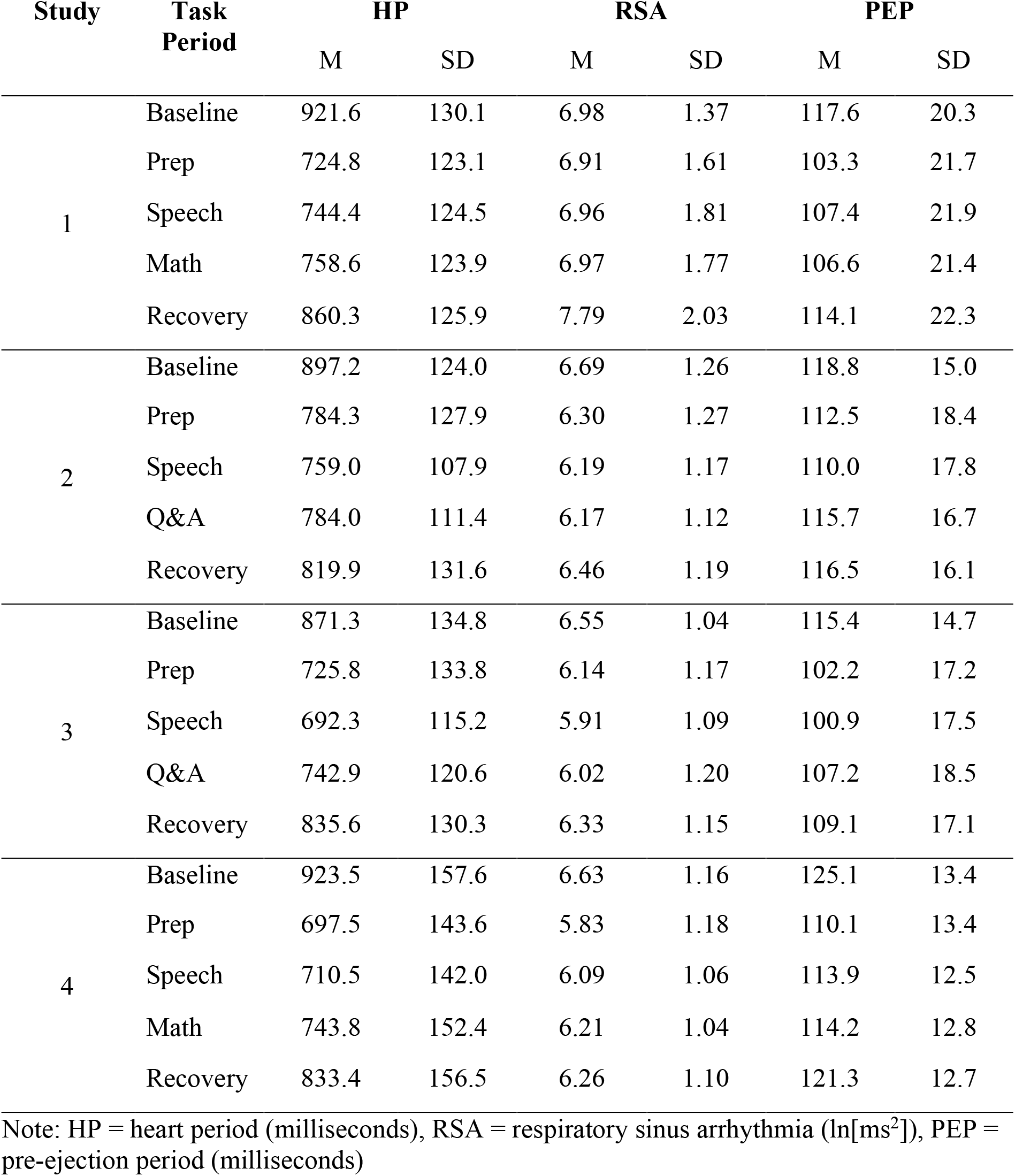
Summary of Autonomic Indices During the Trier Social Stress Test Across 4 Studies.

| Study | Task Period | HP |  | RSA |  | PEP |  |
| --- | --- | --- | --- | --- | --- | --- | --- |
|  |  | M | SD | M | SD | M | SD |
| 1 | Baseline | 921.6 | 130.1 | 6.98 | 1.37 | 117.6 | 20.3 |
|  | Prep | 724.8 | 123.1 | 6.91 | 1.61 | 103.3 | 21.7 |
|  | Speech | 744.4 | 124.5 | 6.96 | 1.81 | 107.4 | 21.9 |
|  | Math | 758.6 | 123.9 | 6.97 | 1.77 | 106.6 | 21.4 |
|  | Recovery | 860.3 | 125.9 | 7.79 | 2.03 | 114.1 | 22.3 |
| 2 | Baseline | 897.2 | 124.0 | 6.69 | 1.26 | 118.8 | 15.0 |
|  | Prep | 784.3 | 127.9 | 6.30 | 1.27 | 112.5 | 18.4 |
|  | Speech | 759.0 | 107.9 | 6.19 | 1.17 | 110.0 | 17.8 |
|  | Q&A | 784.0 | 111.4 | 6.17 | 1.12 | 115.7 | 16.7 |
|  | Recovery | 819.9 | 131.6 | 6.46 | 1.19 | 116.5 | 16.1 |
| 3 | Baseline | 871.3 | 134.8 | 6.55 | 1.04 | 115.4 | 14.7 |
|  | Prep | 725.8 | 133.8 | 6.14 | 1.17 | 102.2 | 17.2 |
|  | Speech | 692.3 | 115.2 | 5.91 | 1.09 | 100.9 | 17.5 |
|  | Q&A | 742.9 | 120.6 | 6.02 | 1.20 | 107.2 | 18.5 |
|  | Recovery | 835.6 | 130.3 | 6.33 | 1.15 | 109.1 | 17.1 |
| 4 | Baseline | 923.5 | 157.6 | 6.63 | 1.16 | 125.1 | 13.4 |
|  | Prep | 697.5 | 143.6 | 5.83 | 1.18 | 110.1 | 13.4 |
|  | Speech | 710.5 | 142.0 | 6.09 | 1.06 | 113.9 | 12.5 |
|  | Math | 743.8 | 152.4 | 6.21 | 1.04 | 114.2 | 12.8 |
|  | Recovery | 833.4 | 156.5 | 6.26 | 1.10 | 121.3 | 12.7 |

Note: HP = heart period (milliseconds), RSA = respiratory sinus arrhythmia ( $\ln[\text{ms}^2]$ ), PEP = pre-ejection period (milliseconds)

At baseline, female participants (*d* = .32, *t* = −*2*.35, *p* = .020), black participants (*d* = .33, *t* = 1.96, *p* = .05), and younger participants (*r* = −.50, *p*<.001) had higher RSA than males, whites, or older participants, respectively. Table 2 summarizes the between-subject associations with RSA and PEP reactivity relative to baseline. Female participants, black participants, and younger participants tended to have greater decreases in RSA during the prep, speech, and math task periods and higher RSA relative to baseline during recovery. Female participants decreased PEP less relative to baseline than male participants during the Q&A task period. Otherwise, no significant differences in PEP reactivity were observed based on sex, race, or age. Individuals with higher baseline RSA had greater decreases in RSA relative to baseline across all the task periods. Individuals with longer PEP at baseline had greater decreases in PEP during the prep, speech, and math task periods but not during Q&A or recovery. The average magnitudes of change in RSA during the prep, speech, and Q&A task periods, but not during math or recovery were significantly correlated with the average magnitudes of change in PEP during the same task periods.

**Table 2:** Between-subject associations with RSA and PEP reactivity.

|  | <u>RSA reactivity</u> |  |  |  |  | <u>PEP reactivity</u> |  |  |  |  |
| --- | --- | --- | --- | --- | --- | --- | --- | --- | --- | --- |
|  | Prep | Speech | Math | Q&A | Recover | Prep | Speech | Math | Q&A | Recover |
|  | Cohen's d |  |  |  |  |  |  |  |  |  |
| Sex (female) | -.43* | -.42* | -.55* | .22 | .26* | .11 | .00 | -.02 | .54* | .17 |
| Race (black) | -.39* | -.39* | -.40* | - | .23 | .05 | -.03 | .06 | - | .17 |
|  | Pearson Correlation |  |  |  |  |  |  |  |  |  |
| Age | .34* | .32* | .32* | - | .29* | .13 | .02 | -.06 | - | .01 |
| Baseline RSA | -.48* | -.49* | -.55* | -.44* | -.31* | .04 | .06 | .07 | .11 | .04 |
| Baseline PEP | .10 | .10 | .10 | .05 | .02 | -.27* | -.23* | -.24* | -.10 | -.09 |
| PEP (same task period) | .17* | .13* | .11 | .34* | .06 | - | - | - | - | - |

Note: RSA = respiratory sinus arrhythmia ( $\ln[\text{ms}^2]$ ), PEP = pre-ejection period (milliseconds), RSA and PEP reactivity are equal to RSA and PEP during that task period minus RSA and PEP at baseline respectively. \* $p < .05$

### RSA-PEP coupling

RSA-PEP coupling at baseline across all participants, regardless of the study or type of feedback received was significant and positive (*β* = .16, *p* < .001). Most of this variance was accounted for by reciprocal responses of RSA and PEP during the prep, speech, math, Q&A, and recovery task periods, but the residual covariance between RSA reactivity and PEP across the 25 1-minute epochs, while small, remained significant (*B*=1.85, *SE* = 0.21, *β*= .04, *p* < .001) (Figure 2).

### Individual differences in RSA-PEP coupling

RSA-PEP coupling was significantly more reciprocal (i.e. positively correlated) for participants with higher baseline RSA in the full sample (Figure 4) and for females among the 258 participants for whose sex had been recorded (Figure 5), but baseline PEP was not significantly associated with RSA-PEP coupling. Females also had significantly higher baseline RSA. The indirect effect of female sex on RSA-PEP coupling via baseline RSA (Figure 5) was significant (*β* = .015; *p* = .028), indicating that more reciprocal RSA-PEP coupling among females is partially due to them having higher baseline RSA. However, female sex still significantly predicted more reciprocal RSA-PEP coupling, accounting for the effects of baseline RSA.

**Figure 4:**
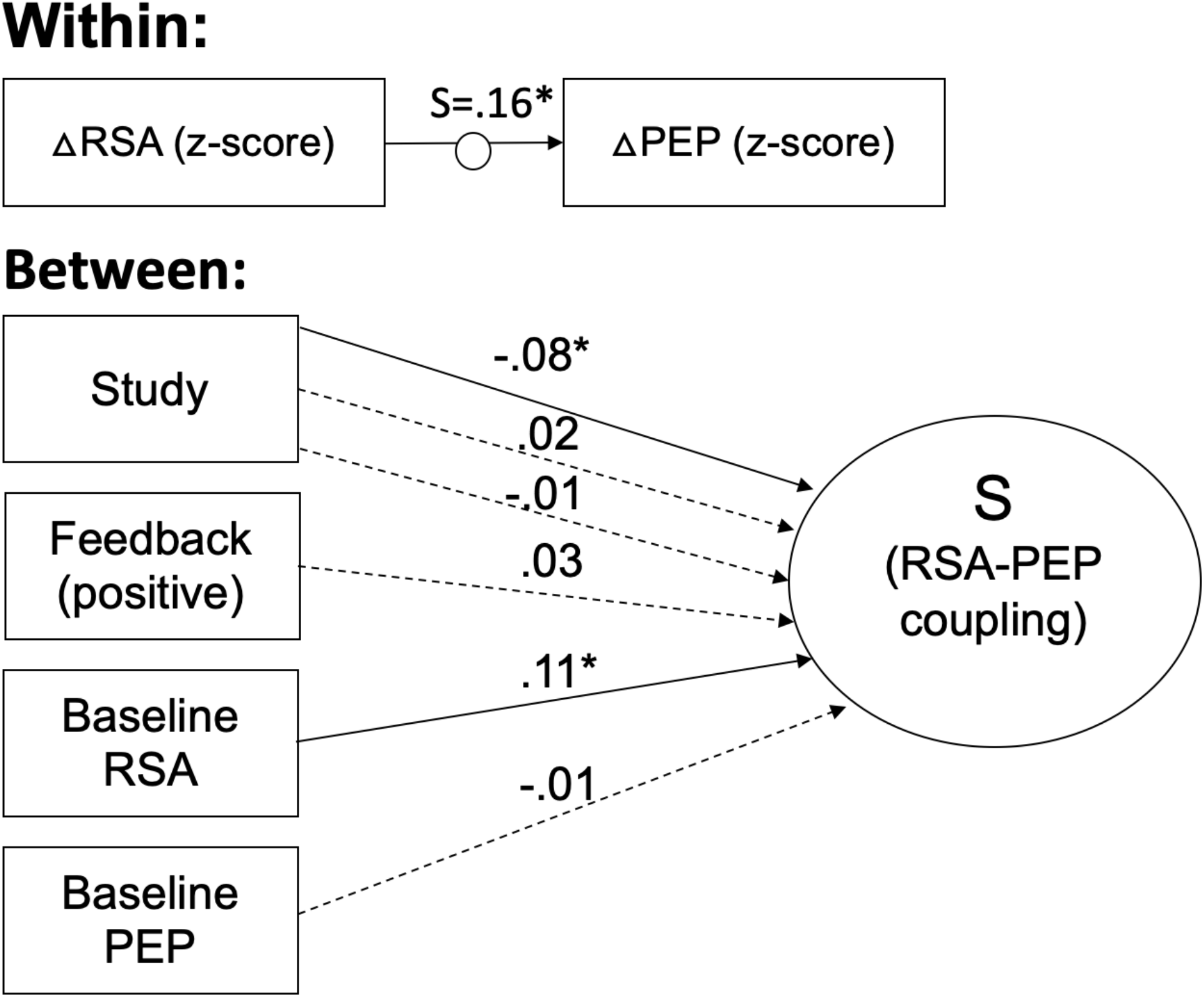
Individual differences in RSA-PEP coupling across 4 studies based on feedback type and baseline RSA and PEP. Results of multilevel structural equation modelling of factors contributing to within-person changes in RSA and PEP (N=325). The change in RSA (ΔRSA) and PEP (ΔPEP) relative to baseline was determined at each of the 25 1-minute epochs of the task and then standardized within subjects. The random slope (S) of the relation between standardized ΔRSA and ΔPEP was used as a between-subjects dependent variable. Coupling during each task period relative to coupling at baseline was evaluated between subjects. Between subjects, “Study” refers to categorical variables for the 4 studies from which data was combined for this analysis. “Feedback” represents whether, for that participant, confederate judges responded with positive (i.e. smiling) compared to with the traditional neutral expression. Coefficients are standardized. Solid lines indicate significant paths. Dotted lines indicate nonsignificant paths. *p < .05

**Figure 5:**
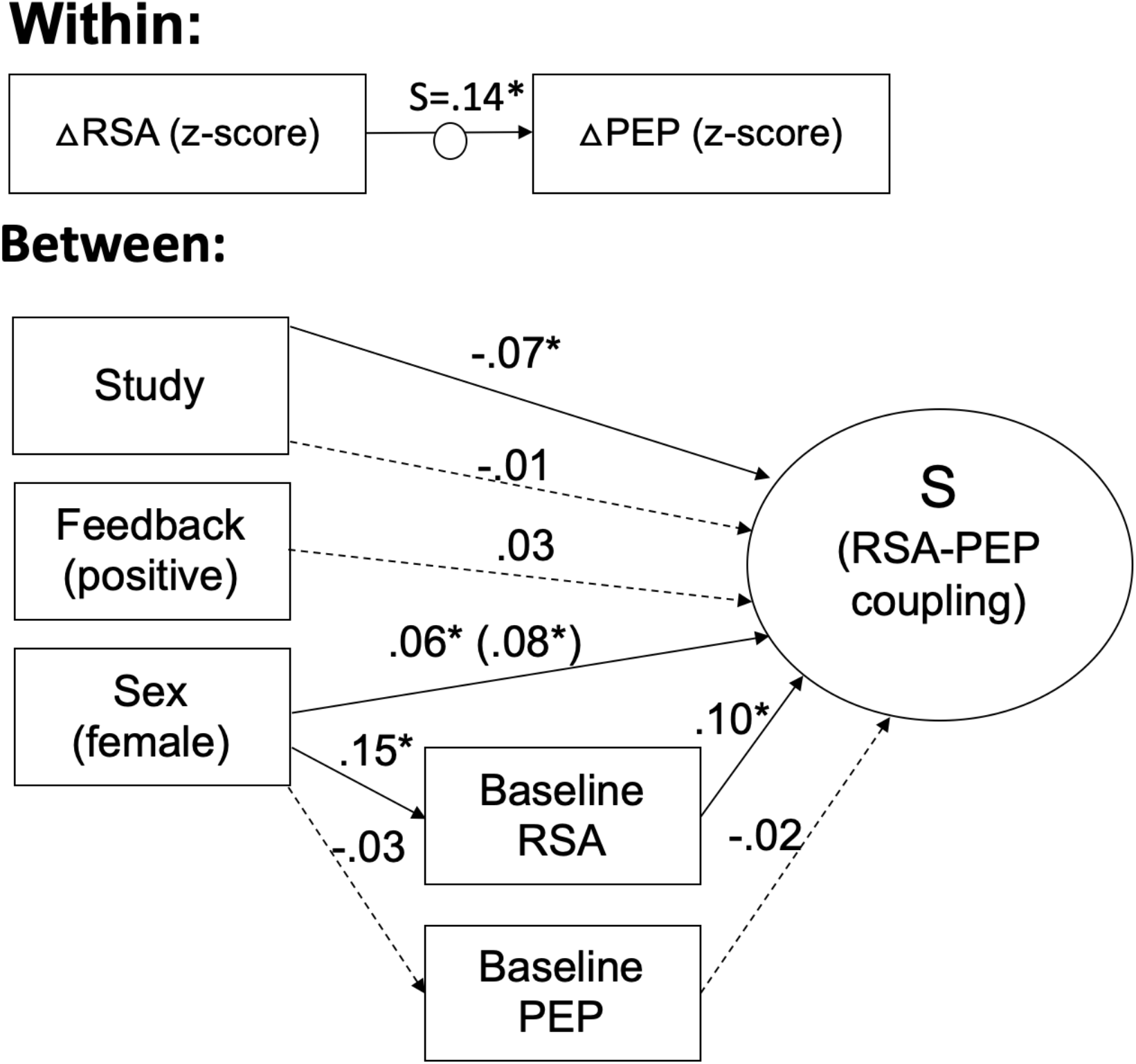
Individual differences in RSA-PEP coupling across 3 studies based on feedback type, baseline RSA and PEP, and sex. Results of multilevel structural equation modelling of factors contributing to within-person changes in RSA and PEP (N=258). The change in RSA (ΔRSA) and PEP (ΔPEP) relative to baseline was determined at each of the 25 1-minute epochs of the task and then standardized within subjects. The random slope (S) of the relation between standardized ΔRSA and ΔPEP was used as a between-subjects dependent variable. Coupling during each task period relative to coupling at baseline was evaluated between subjects. Between subjects, “Study” refers to categorical variables for the 3 studies from which data was combined for this analysis. Participants in the fourth study were excluded because the sex of the participants was not recorded. “Feedback” represents whether, for that participant, confederate judges responded with positive (i.e. smiling) compared to with the traditional neutral expression. Coefficients are standardized. Solid lines indicate significant paths. Dotted lines indicate nonsignificant paths. *p < .05

Among the 131 participants for whom information on age, race, and sex were available, coupling was significantly more reciprocal for females, younger participants, and participants with higher baseline RSA (Figure 6). The association between age and RSA-PEP coupling was fully mediated by younger participants having higher baseline RSA (Indirect Effect: *β* = −.044; *p* = .011). The associations between race and RSA-PEP coupling and between race and baseline RSA were not significant when accounting for the effects of age, sex, and baseline ANS activity.

**Figure 6:**
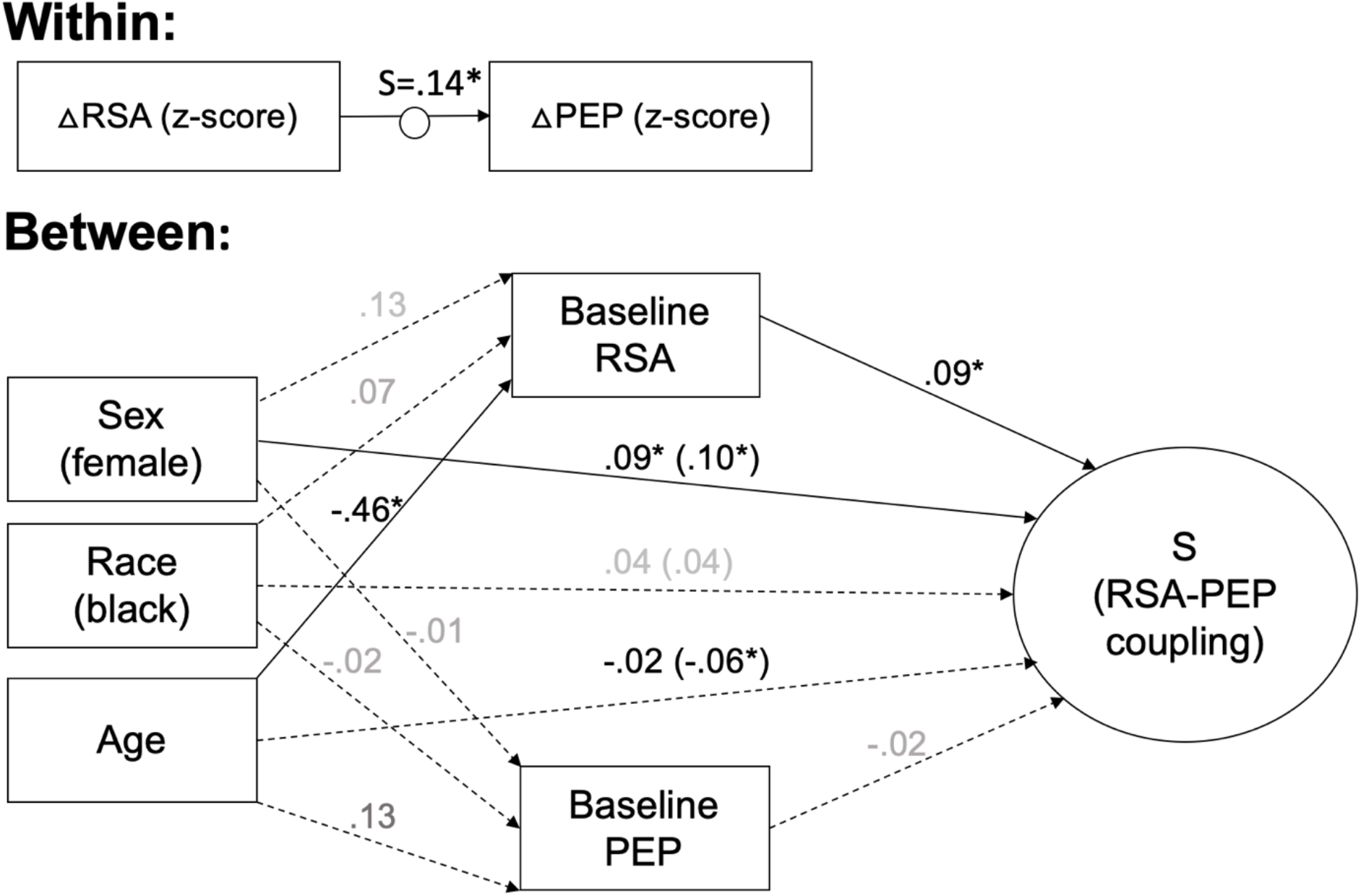
Individual differences in RSA-PEP coupling based on sex, race, age, and baseline autonomic activity. The change in RSA and PEP relative to baseline during each task period was assessed within subjects. The random slope (S) of the relation between RSA and PEP, standardized within-subjects was used as a between-subjects dependent variable. The random slope (S) of the relation between RSA and PEP, standardized within-subjects was used as a between-subjects dependent variable. Coupling during each task period relative to coupling at baseline was evaluated between subjects. Coupling during each task period relative to coupling at baseline was evaluated between subjects. Coefficients are standardized. Solid lines indicate significant paths. Dotted lines indicate nonsignificant paths. *p < .05

## Discussion

This study investigated the nature of variability and flexibility in SNS-PNS coupling between and within individuals at rest, and in response to an acute stressor. The results indicate that the SNS and PNS are reciprocally coupled with one another at rest and in response to stressors. Increases in SNS activity tend to be associated with decreases in PNS activity. This reciprocal activity is due to both opposite response patterns to acute stress, but also exist within-situations (e.g. task periods) in which stress exposure is comparable, perhaps reflecting shared control mechanisms in the central nervous system. Conversely, certain states, in particular recovery following an acute stressor, seem to be characterized by coactivation. On average, RSA was higher than baseline, and PEP was shorter than baseline during the recovery period, indicating increased activation of both the SNS and PNS relative to baseline during recovery. Moreover, individuals differ in the strength of their SNS-PNS coupling. Specifically, females and those with higher baseline RSA, in particular younger participants, have more reciprocal SNS-PNS coupling across the TSST.

Across four studies, the SNS and PNS were reciprocally coupled, such that increases in SNS activity tend to be associated with decreases in PNS activity. This is evidenced by the significant positive between-person correlations between the magnitude of RSA and PEP reactivity to a standardized laboratory stressor, and the significant positive within-person relation between PEP and RSA across the TSST, even accounting for differences in PEP and RSA between the parts of the task. These reciprocal associations are consistent with prior findings (Cacioppo et al., 1994a) and may reflect shared mechanisms of autonomic regulation in the brain. Regions in the brain’s central autonomic network have been found to regulate both the PNS and the SNS. In particular, amygdala activation leads to increases in SNS activity and decreases in PNS activity through activation or disinhibition of sympathoexcitatory neurons in the rostral ventrolateral medulla, and inhibition of vagus nerve activity through the nucleus ambiguus (Thayer & Lane, 2009), and activity in the ventromedial prefrontal cortex has been found to have an inhibitory influence on the amygdala and to lead to increases in PNS activity and decreases in SNS activity (Thayer & Lane, 2009; Zhang et al., 2014). Therefore, while we do find that RSA-PEP coupling is varies depending on the environmental demands, as predicted by the autonomic space model (Berntson et al., 1991), we also find evidence of a tendency toward reciprocal activation within and across task periods, perhaps reflecting these shared mechanisms of regulation independent of the response to a stressor.

During recovery from an acute stressor RSA was significantly higher than baseline on average, whereas PEP was significantly shorter, suggesting increased activation in both the SNS and PNS relative to baseline. Taken together, these results suggest that while SNS activity may be slow to return to baseline following a stressor, PNS activity may be augmented to promote recovery, consistent with previous observations of “vagal rebound” (Mezzacappa et al., 2001; Page-Gould et al., 2010). This suggests that certain environmental demands may be associated with non-reciprocal responses from the two autonomic branches.

We also observed that individuals differed in the extent to which SNS and PNS activity were reciprocally coupled across the TSST. Females, younger adults, and individuals with higher baseline RSA have more reciprocally coupled SNS and PNS activity. Females and younger adults also had higher baseline RSA on average. Sex but not age differences in RSA-PEP coupling remained significant when accounting for the relation with baseline RSA. Individuals with higher baseline RSA are hypothesized by polyvagal theory to have more flexibility in emotional responses due to a greater range of potential physiological states (Porges et al., 1994). Indeed, higher baseline RSA has been shown to be related to greater decreases in RSA in response to social stressors (Rigoni et al., 2017), as would be expected by the law of initial values (Berntson et al., 1991). This may lead to more reciprocal coupling, as increases in SNS activity indicated by shorter PEP may be accompanied by greater reciprocal PNS withdrawal in individuals with higher baseline RSA. Similarly, the combination of higher baseline RSA and more reciprocal coupling for females presents a physiological profile with a broader range of physiological reactivity, but also a more consistent, reciprocal relationship between the SNS and PNS.

### Limitations and Future Directions

While this study has many strengths, including a large sample size combined across several studies using the same acute stressor, it does have some limitations that raise outstanding questions for future research. First, because of inconsistency in demographic, mental health, and other individual difference variables collected across the studies, we were limited to sex and baseline autonomic activity as indicators of potential individual differences in multiple studies. Our power was therefore limited to detect possible age and race-related differences in RSA-PEP coupling. Future work examining RSA-PEP coupling with larger samples and in relation to characteristics of emotion functioning like mental health and emotion regulation could further illuminate how the extent to which SNS and PNS activity are correlated and may vary between individuals and contribute to emotion functioning. Further, while the studies included all used the TSST, protocols were similar, and differences were controlled for statistically, inconsistencies nonetheless existed. For example, a Q&A period replaced the math task in studies that used positive feedback because mental arithmetic typically results in a diverting of attention away from the evaluators (looking away, closed eyes). Replication in large sample size studies with consistent protocols could assess the generalizability and replicability of the findings of these analyses. In addition, the length of the epochs used here to reflect coupling likely impacts the results to some extent, which may make it more difficult to generalize these results to future work that uses different epoch lengths. However, 1 minute epochs are common and reliable units of analyses in research on autonomic functioning, and using shorter epochs has costs for the reliability of the measurement of RSA especially (Beauchaine et al., 2019; Berntson et al., 1997). Future work should nonetheless examine the temporal dynamics of autonomic control at different levels of temporal precision. Finally, response to and recovery from an acute social stressor is only a small slice of the complex social and emotional milieu that individuals navigate daily. Future work should evaluate SNS-PNS coupling across a broader range of social and emotional situations and inductions.

## Conclusion

Through a reanalysis of data from four different studies, we present a coherent picture of the correlated activity of the PNS and SNS at baseline and in response to an acute stressor. Overall, the SNS and PNS respond inversely to stress and are reciprocally coupled, perhaps reflecting shared regulatory mechanisms in the brain and reciprocal responses to environmental demands. However, recovery from a stressor is characterized by coactivation on average as the SNS recovers slowly, while PNS activity rebounds and augments to accelerate recovery. Individuals also vary in the extent to which their SNS and PNS are reciprocally coupled, with females, younger adults, and individuals with higher baseline RSA demonstrating more reciprocal coupling, reflecting a more coherent pattern of autonomically-mediated physiological responding.

## Acknowledgments

This work was supported by National Institutes of Health grant 5T32MH020006-19 (Johnson, Gross) and an NIA grant R24AG048024 (Epel, Mendes).

